# Identifying Spatial Co-occurrence in Healthy and InflAmed tissues (ISCHIA)

**DOI:** 10.1101/2023.02.13.526554

**Authors:** Atefeh Lafzi, Costanza Borrelli, Karsten Bach, Jonas A. Kretz, Kristina Handler, Daniel Regan-Komito, Xenia Ficht, Andreas Frei, Andreas Moor

## Abstract

Spatial transcriptomics techniques are able to chart the distribution and localization of cell types and RNA molecules across a tissue. Here, we generated matched sequencing-based (Visium) and hybridization-based (Molecular Cartography) spatial transcriptomics data of human IBD samples. We then developed ISCHIA (Identifying Spatial Co-occurrence in Healthy and InflAmed tissues), a computational framework to analyze the spatial co-occurrence of cell types and transcript species in the tissue environment. ISCHIA revealed tightly associated cellular networks, ligand-receptor interactions enriched in the inflamed human colon, and their associated gene signatures, highlighting the hypothesis-generating power of co-occurrence analysis on spatial transcriptomics data.

## Background

Tissue ecosystems are maintained by the co-existence and coordinated function of cellular networks (CNs). CNs are the basic functional unit of any tissue: communities of neighboring cells that interact to perform a physiological function, with cells as nodes and cell-cell interactions (CCIs) as edges. A prime example of CN is the colonic crypt, where a network of Wnt-secreting mesenchymal cells and Wnt-receiving epithelial stem cells provides self-renewal and regenerative capacity to the colonic epithelium^1^.

During inflammation, CNs can be perturbed by the induction of aberrant cell states and recruitment of non-resident cell types, much like natural ecosystems are disturbed by the arrival of alien species^2^. Inflammatory CNs can facilitate tissue regeneration and reestablishment of homeostasis, or can act as drivers of pathology, especially in chronic settings such as inflammatory bowel diseases (IBD). Understanding how CN architecture is altered by inflammation is fundamental to gaining comprehensive insights into pathological mechanisms. Most recently, CNs have been inferred from single-cell RNA-Sequencing (scRNASeq) data by computationally predicting CCIs based on the expression levels of ligands and their receptors^3,4^. Due to tissue dissociation, scRNASeq provides a fragmented view of a tissue ecosystem: cells predicted to interact might populate spatially distinct areas of the tissue, and thus are unlikely to constitute a genuine CN. Hence, there is a need for analytic methods that leverage spatial information to shortlist and prioritize CCIs that are more likely to occur in intact tissues. Next generation sequencing (NGS)-based spatial transcriptomics (ST) methods, such as Visium ST (10X Genomics), capture RNA molecules *in situ* at spatially barcoded spots, generating bulk RNA profiles of 10-30 cells^5^. While this is generally considered a disadvantage, we show here that these “mixed transcriptomes” can be used to infer CNs, as their gene expression profiles contain information about 1) cell type composition, 2) expressed ligand receptor (LR) pairs, and 3) signaling pathways, gene regulatory networks and effector molecules that mediate CN function. We hypothesize that the inherent spatial proximity of spot data can be leveraged to integrate these three levels of information, reconstructing the architecture and function of CNs. To this end, we developed ISCHIA (Identifying Spatial Co-occurrence in Healthy and InflAmed tissues), a computational framework that assigns a quantitative property to the interaction potential of cell types or LR pairs by computing their spatial co-occurrence. This probabilistic approach was inspired by species co-occurrence models in ecology, which derive statistically significant patterns of pairwise species associations from the frequency of their observed co-occurrence at defined spatial locations^6^. Co-occurrence between two species may be positive (observed co-occurrence is higher than expected by chance), random (independently distributed), or negative (lower than expected by chance). For decades, ecologists have analyzed co-occurrence patterns of plant and animal species to understand ecological communities and the rules of their assembly ^6^. Here, we apply the same principles to gain deeper insights into cellular communities and the rules of their spatial associations in tissue biology.

We applied ISCHIA to chart the CN landscape of the healthy and inflamed human colon. We generated both sequencing-based (Visium) and hybridization based (Molecular Cartography) data of human ulcerative colitis samples. First, we applied ISCHIA to identify pairs of cell types that are positively co-occurring in inflammation-specific cellular neighborhoods. Within these neighborhoods, ISCHIA then performs co-occurrence analysis of ligand and receptor genes, and derives the associated gene signatures. Thereby, we reconstructed the architecture of CNs, such as a M cell-fibroblast network enriched in the inflamed colon. Next, we extended co-occurrence analysis to the whole interactome (all ligands and receptor genes) to uncover spatially coordinated tissue responses to inflammation. Finally, we applied co-occurrence analysis to a spatial transcriptomics dataset of murine colitis and identified conserved tissue repair pathways.

## Results

### Composition-based clustering of spots divides tissues into composition classes

We generated Visium data from inflamed and non-inflamed colon regions of 4 ulcerative colitis (UC) samples (**Supplemental Fig S1a**). As inferring CNs in each and every spot separately would be noisy, computationally intensive, and would lack statistical power, ISCHIA first divides the tissue into clusters of similar cellular composition - termed composition classes (CCs) (**Fig 1a**). CCs are thus groups of spots containing similar cellular communities, e.g, all spots capturing colonic crypts. To achieve the division of the tissue into CCs, spot transcriptomes are deconvoluted, yielding a cell type composition matrix (spot × contribution of each cell type), which is then subjected to dimensionality reduction and k-means clustering. ISCHIA allows for both reference-free and reference-based deconvolution. Here, we took advantage of published^7–9^ and in-house datasets to compile a comprehensive, integrated IBD scRNAseq reference (**Supplemental Fig S1b**), which was used for spot deconvolution (see Methods). Notably, the Handler *et al*.^9^ dataset was collected using the microwell-based platform BD Rhapsody, and thus contains granulocytes that are mostly absent from data generated with droplet-based methods.

**Figure 1.**
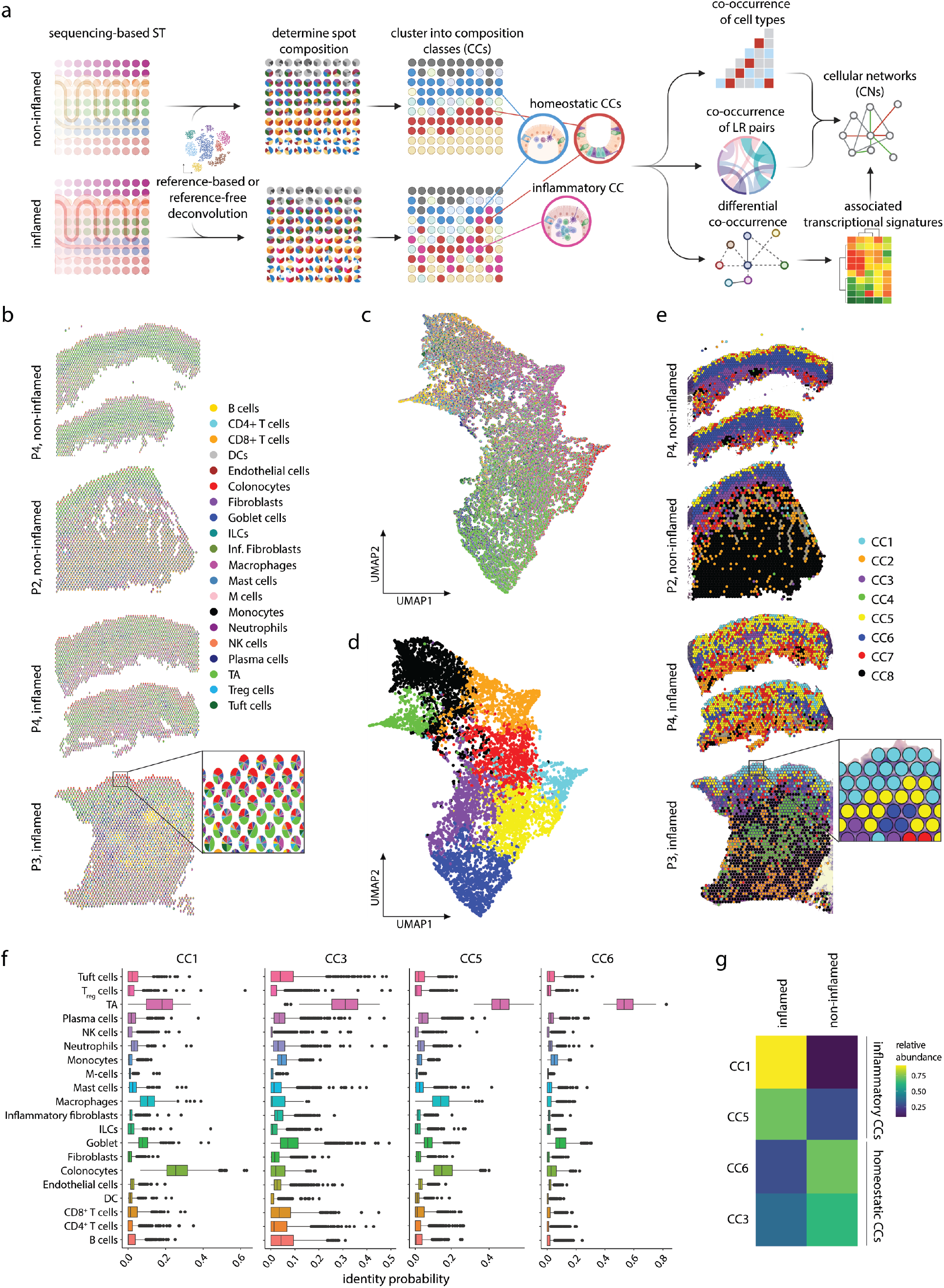
ISCHIA performs composition-based clustering of spot data. **a,** Schematic workflow of the ISCHIA pipeline. **b,** Deconvolution of Visium spots based on IBD scRNASeq reference data. Spot composition is visualized as a pie chart (shown in inset) and projected on the spatial coordinates. **c,** Dimensionality reduction of deconvoluted matrix from all four UC samples. **d,** Composition-based k-means clustering of the deconvoluted matrix yields 8 composition classes (CC1-8) reflecting similar cell type compositions. **e,** Spots overlayed on tissue coordinates colored by CC. **f,** Predicted cell-type proportions in CCs. **g,** Relative abundance of CCs in inflamed and non-inflamed samples. Abbreviations: CC, composition class; TA, transit amplifiers; ILC, innate lymphoid cells; DCs, dendritic cells.

We performed joint deconvolution of all spots across all samples (**Fig 1b**), followed by k-means clustering of the deconvoluted matrix (**Fig 1c**), revealing 8 CCs of co-localizing cell types (**Fig 1d**). In the healthy colon, CCs broadly reflect component layers of the colonic wall: submucosa (CC3), crypt bottom (CC6), and crypt top (CC5 and CC1) **(Fig 1e**). In the inflamed samples, the spatial boundaries between CCs are lost, indicating perturbed tissue architecture and altered spatial arrangement of cells **(Fig 1e**). Of note, CC2, CC4, CC7 and CC8 were not considered for downstream analysis as they mapped onto the muscular layer or were highly patient-specific (**Supplemental Fig S1c**). The most prevalent cell types across all composition classes are transit amplifiers (TAs), colonocytes and goblet cells, reflecting the cellular composition of the adult colonic epithelium^10^ (**Fig 1f**). CC1 and CC5 are enriched in the inflamed colon and were therefore termed inflammatory CCs (**Fig 1g**). Indeed, their cellular composition reflects the recruitment of macrophages, and hallmark of IBD pathogenesis^11,12^.

These results indicate that subdividing tissue into CCs not only recapitulates morphology and architecture but is also able to capture alterations in cell type composition across conditions. Most importantly, this composition-based clustering of spots can be used to focus downstream differential gene expression analysis to specific groups of cells, effectively filtering out transcriptome heterogeneity arising from distinct cellular compositions.

### Cellular co-occurrence in the inflamed colon

We hypothesized that inflammatory CCs (CC1 and CC5) would contain information about non-homeostatic CNs induced by spatial rearrangement of cells and infiltrating leukocytes. We thus applied co-occurrence analysis of cell types to inflammatory and the homeostatic CCs of both inflamed and non-inflamed colon samples. ISCHIA identified several pairwise cell type co-occurrences as being significantly more frequent than expected by chance in inflammatory CCs (positive co-occurrence, *P < 0.05*) (**Fig 2a, b**). For example, many cellular co-occurrences involving M-cells - highly specialized epithelial cells involved in antigen presentation^13^ - were positively co-occurring in inflammatory CCs but not in homeostatic CCs. Of note, ISCHIA’s co-occurrence predictions are irrespective of cell type abundance. Indeed, being the most prominent cell type in all CCs, TAs are observed to be co-occurring with many cell types. However, this is not higher than what is expected given their abundance.

**Figure 2.**
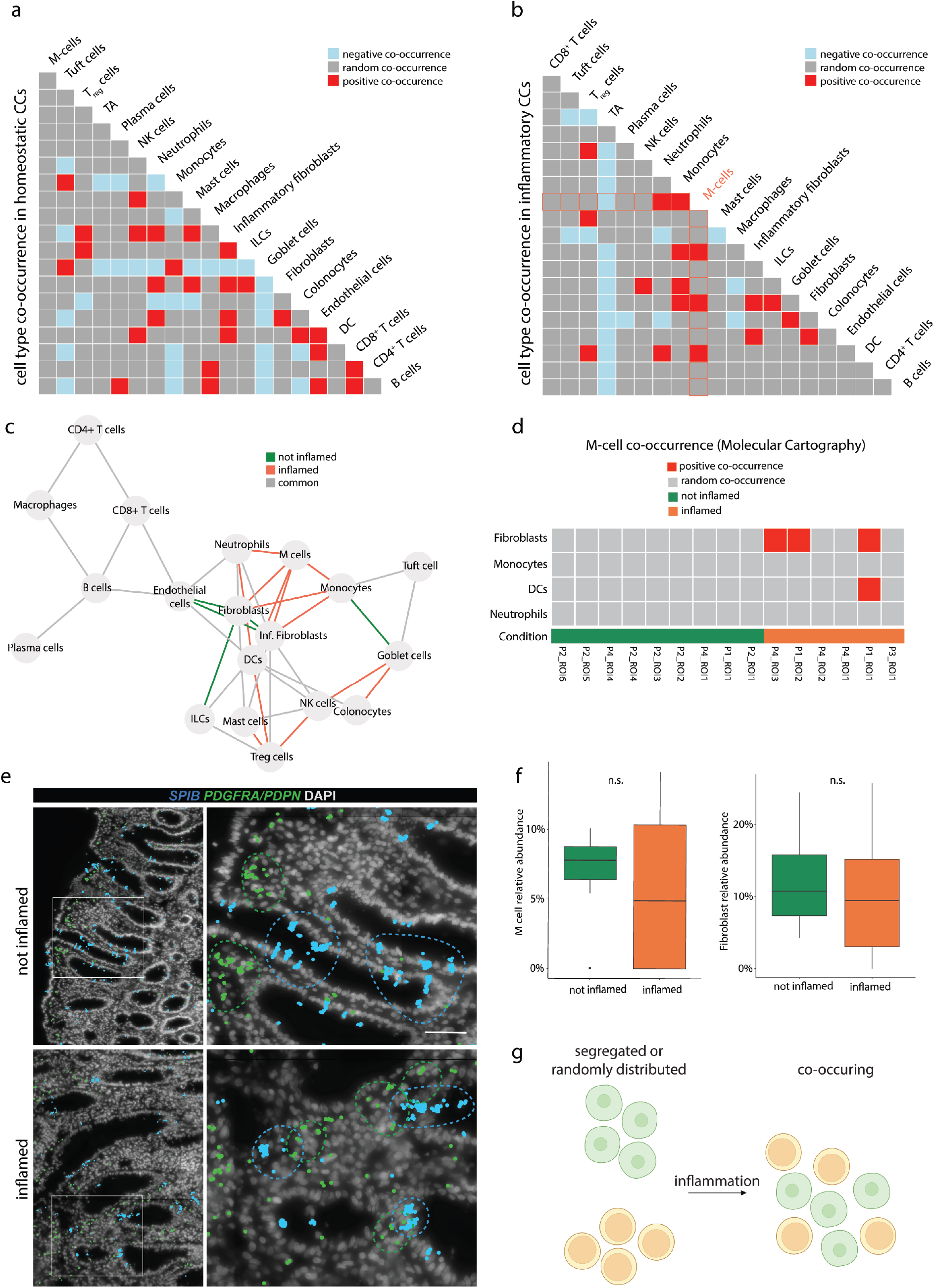
M-cells and fibroblasts co-occur in the inflamed human colon. **a, b,** Diagonal matrix plot depicting cell type co-occurrences in homeostatic CCs (left) and inflammatory CCs (right). Co-occurrence is positive when observed more frequently than expected *(P < 0.05)*, random when there is no significant difference, negative when observed less than expected *(P < 0.05)*. **c**, Cell type co-occurrences in inflammatory CCs, shown as a network. Nodes are cell types, edges indicate positive co-occurrences, colored by condition. **d,** M-cell co-occurrence in spatially restricted cellular neighborhoods (k = 5) quantified separately for each ROI from Molecular Cartography data. **e,** *SPIB* (M-cells) and *PDGFRA/PDPN* (fibroblasts and inflammatory fibroblasts) expression in inflamed and non-inflamed colon as shown by Molecular Cartography. Dashed lines indicate areas of elevated SPIB (blue lines) or PDGFRA/PDPN (green lines) expression. Scale bar 20 μm. **f,** M-cell *(SPIB+* segments) and fibroblasts *(PDGFRA/PDPN+* segments) relative abundance, as observed in 6 UC patient samples by Molecular Cartography. n.s., non significant, unpaired Wilcoxon test. **g,** Spatial rearrangement of cells during inflammation leads to positive co-occurrence.

We further subdivided spots of inflammatory CCs in those coming from inflamed vs non-inflamed samples, and found that M-cell co-occurrence were specifically induced by inflammation (**Fig 2c**). Recently, M-like cells were indicated as an interaction hub during colitis based on CCI predictions from scRNASeq data^7^. Here, we complement this finding with spatial information and prioritize M-cell interactions with monocytes, neutrophils, dendritic cells, and fibroblasts based on the co-occurrence of these cell types specifically in the inflamed colon.

To validate ISCHIA’s co-occurrence predictions from spot data (bulk) on a single cell level, we performed 100-plex RNA fluorescent *in situ* hybridization (FISH, Molecular Cartography). Contrarily to sequencing-based ST methods such as Visium, multiplexed *in situ* hybridization techniques such as Molecular Cartography achieve singlecell resolution. Co-occurrence can thus be calculated in even smaller cellular neighborhoods (e.g., k nearest neighbors = 5 cells), greatly restricting the interaction space and further refining predictions. For this analysis, we included an additional inflamed and non-inflamed sample to the 4 samples subjected to Visium, for a total of 6 human UC colon samples. Nuclear DAPI staining was used to segment Molecular Cartography data into putative single cells. Segments were then annotated based on marker gene expression and subsequently used as input for cell type co-occurrence analysis. Confirming predictions by ISCHIA based on spot data, Molecular Cartography revealed that M-cells (*SPIB*+ segments) and fibroblasts (*PDGFA/PDPN+* segments) were significantly co-occurring in inflamed but not in non-inflamed samples (**Fig 2d, e**). Importantly, the increased M-cell and fibroblast co-occurrence was independent of their abundance (**Fig 2f**). Thus, ISCHIA has the capacity to capture interactions arising through co-occurrence that has been newly established - possibly by spatial rearrangement or altered distribution of the involved cell types - even if cell type frequencies are unaltered (**Fig 2g**).

Co-occurrence analysis can therefore be used on spot data, as well as FISH-based ST data, to infer interaction potential of cell types based on their proximity. In ecology, co-occurrence is not generally considered evidence of interaction^14^, however, we argue that it can be used as a measure of spatial association and proximity, prerequisites for juxtacrine and paracrine signaling between cells.

### Linking cell type and ligand-receptor co-occurrence to reconstruct cellular networks

We next applied spatial co-occurrence analysis to transcript species encoding for ligand and receptor genes. Indeed, we reasoned that the potential for molecular interaction (juxtacrine and paracrine signaling) is higher between positively co-occurring LR pairs, that is pairs that co-occur in the tissue more frequently than expected by chance. To test this hypothesis, we performed co-occurrence analysis of LR genes within one of the inflammatory CCs, CC5, and identified significantly positively co-occurring pairs (**Supplementary Table 1**). We next ranked them based on the correlation of their expression within CC5 spots (see Methods, top 20 shown in **Fig 3a**). Several of the significantly co-occurring LR pairs were previously implicated in colitis, and are mostly involved in epithelial integrity and repair (*DDR1-CDH1*^15^, *DSG2*^16^), immune recruitment (*MIF-CD74*^17^), and TNF signaling (*GRN-TNFRSF1A*^18,19^). This highlights ISCHIA’s ability to capture disease-relevant pathways from spatial transcriptomics data, even at a limited sample size.

**Figure 3.**
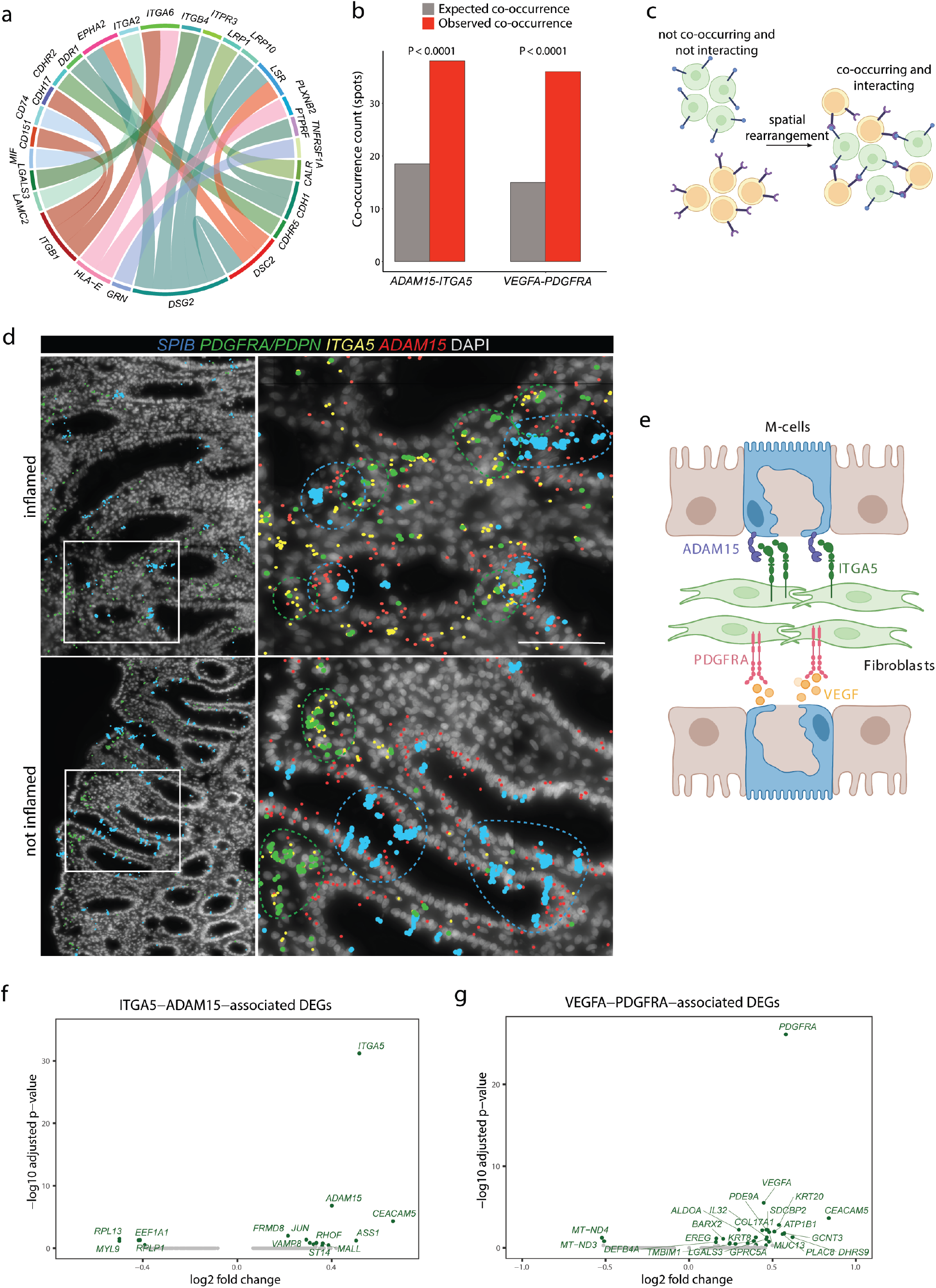
Reconstructing cellular networks from cell type co-occurrence, LR co-occurrence and corresponding transcriptomic signatures. **a**, Top 20 positively co-occurring LR pairs in inflammatory CC5 (observed co-occurrence > expected co-occurrence). Ranking based on correlation of expression of ligand and receptor genes within spots**. b,** Expected vs observed co-occurrence of *ADAM15* and *ITGA5* and *VEGFA* and *PDGFRA*. **c,** Spatial rearrangement of cells induces positive co-occurrence of LR pairs even if expression levels are unaltered. **d,** *SPIB, PDGFRA/PDPN, ADAM15* and *ITGA5* expression in inflamed and non-inflamed colon as shown by Molecular Cartography. Dashed lines indicate areas of elevated SPIB (blue lines) or PDGFRA/PDPN (green lines) expression. Scale bar 20 μm. **e,** Proposed cellular network in inflammatory CCs. **f, g** DEGs associated with ITGA5-ADAM15 and VEGFA-PDGFRA pairs.

To test the hypothesis that positively co-occurring cell types potentially interact via positively co-occurring LR pairs, we focused on M-cells and fibroblasts. Among positively co-occurring LR pairs in inflammatory CC5, we identified two ligands expressed by M cells *(ADAM15* and *VEGFA)* in our integrated IBD scRNASeq reference, and their respective receptors expressed by fibroblasts *(ITGA5* and *PDGFRA)* (**Fig 3b, Supplemental Fig S2a**). Importantly, ISCHIA’s predictions are agnostic to gene expression levels (gene count threshold ≥ 1) and only take into account the proximity (measured as co-occurrence) of a given ligand and receptor (see Methods). As for cell types, ISCHIA identifies interactions arising through positive co-occurrence of LR pairs. This reveals interactions that specifically occur in inflamed samples, even if the expression of the encoding LR genes are unaltered compared to non-inflamed samples, implying a spatial rearrangement of the LR-expressing cells (**Fig 3c**). Indeed, while the co-occurrences of *ITGA5* and *ADAM15*, and *VEGFA* and *PDGFRA* are positive in inflammatory CC5, their expression within these CC is unaltered (**Supplemental Fig S2b**).

Molecular Cartography further confirmed the expression of *ITGA5* by *PDGFA/PDPN+* fibroblasts and of *ADAM15* by *SPIB+* M cells (**Fig 3d**). By linking cell type and LR co-occurrence we thus identify a new CN arising in inflammation (**Fig 3e**). Interestingly, an independent immunohistochemical analysis of IBD colon sections showed close contact between ADAM15-positive epithelial cells and α5β1(*ITGA5*)-positive myofibroblasts in regenerative areas^20^. While functional characterization of this interaction in IBD is outstanding, it illustrates the hypothesis-generating power of co-occurrence analysis performed by ISCHIA.

### Integrating cell types, LR pairs, and transcriptomic signatures to infer CN function

We reasoned that an LR pair is expected to be associated with a specific transcriptional signature, which may be either upstream (triggering LR expression) or downstream of the interaction (consequences of LR signaling). As spot data are only temporal snapshots, we cannot distinguish between causative and consequential effects, but capture the sum of both as a LR-associated transcriptional signature.

After identifying putatively disease-relevant LR pairs, ISCHIA performs differential gene expression analysis between spots of the same CC expressing or lacking that LR pair, in order to uncover the associated transcriptional responses. Importantly, by doing this within the same CC, we filter out transcriptomic effects arising from differences in cell type composition, and only capture effects derived from alterations in cell state that are associated with the presence or absence of a given LR pair. Within CC5, we computed differentially expressed genes (DEGs) between spots having or lacking *ITGA5-ADAM15* expression (**Fig 3f**). Pathway analysis of the resulting significant DEGs revealed enrichment of terms related to fibronectin matrix formation as well as integrin-linked kinase signaling and signaling events mediated by focal adhesion kinase (FAK) (**Supplemental Fig. S2c**). These results corroborate recent studies indicating that *ITGA5* is important for fibronectin assembly by fibroblasts^21^ and that ADAM15 interaction with integrin αV *(ITGA5)* activates FAK signaling to promote migration^22^. Of note, these pathways were not enriched in *VEGFA-PDGFRA*-associated gene sets, which instead are involved in cell-junction organization and the HIF-1α transcription factor network (**Fig 3g, Supplemental Fig. S2d**), possibly indicating a coordinated hypoxia-response.

Using spot data from unbiased, genome-wide spatial transcriptomic profiling, presence-absence information (cell type composition, cell type co-occurrence, and LR co-occurrence) can thus be correlated to alterations in the transcriptome, granting deeper insights into the function of reconstructed CNs.

### Differential co-occurrence identifies niche-specific response programs

Ligands and receptors genes expressed within a spot make up the words of the locally occurring cellular ‘conversations’. We thus computed the co-occurrence of all ligands and receptors (i.e. interactome components) within all spots, irrespective of whether they are annotated as interacting pairs based on PPI predictions databases. We next selected differentially co-occurring interactome components: pairs of ligands and receptor genes that only positively co-occur in a composition class *(P_CCx_ < 0.05 & P_CCy_ > 0.05*) or in a condition *(P_condition_x_ < 0.05 & P_condition_y_ > 0.05*). We reasoned that these particular ‘conversations’ may reveal coordinated tissue responses specific to distinct niches or disease-relevant states. interactome components unique to the inflammatory CC5, which is characterized by a high relative abundance of macrophages. We identified 482 differentially co-occurring, pair-wise interactome combinations that are significantly positively co-occurring in CC5 (*P_CC5_ < 0.05)* but not in other CCs *(P_other_CCs_ > 0.05)*, across both inflamed and not-inflamed samples, indicating their tight association with this particular cell type composition (**Fig 4a**). Among those, we recovered predicted protein-protein interactions such as *BMP2-FGFR2, APP-BDKRB1* and *EDN1-F2RL1* (from the NicheNet database). The strongest differentially co-occurring LR network (ranked by P value difference and observed co-occurrence in CC5) involve *EDN1* - an endothelium-derived vasoconstricting peptide implicated in the pathogenesis of IBD^23^. We also detected differential co-occurrence of the epithelium-derived pro-angiogenic factors *SEMA3C^24,25^* and *CXCL5^26^*. Molecular Cartography confirms localized expression of epithelial *SEMA3C* with *PTPRB* (expressed by the endothelium), *MMP9* (macrophages and monocytes), *CXCL8* (macrophages), *CXCL5* (epithelium, inflammatory fibroblasts, neutrophils) and *SEMA4D* (T cells) (**Fig 4b**).

**Figure 4.**
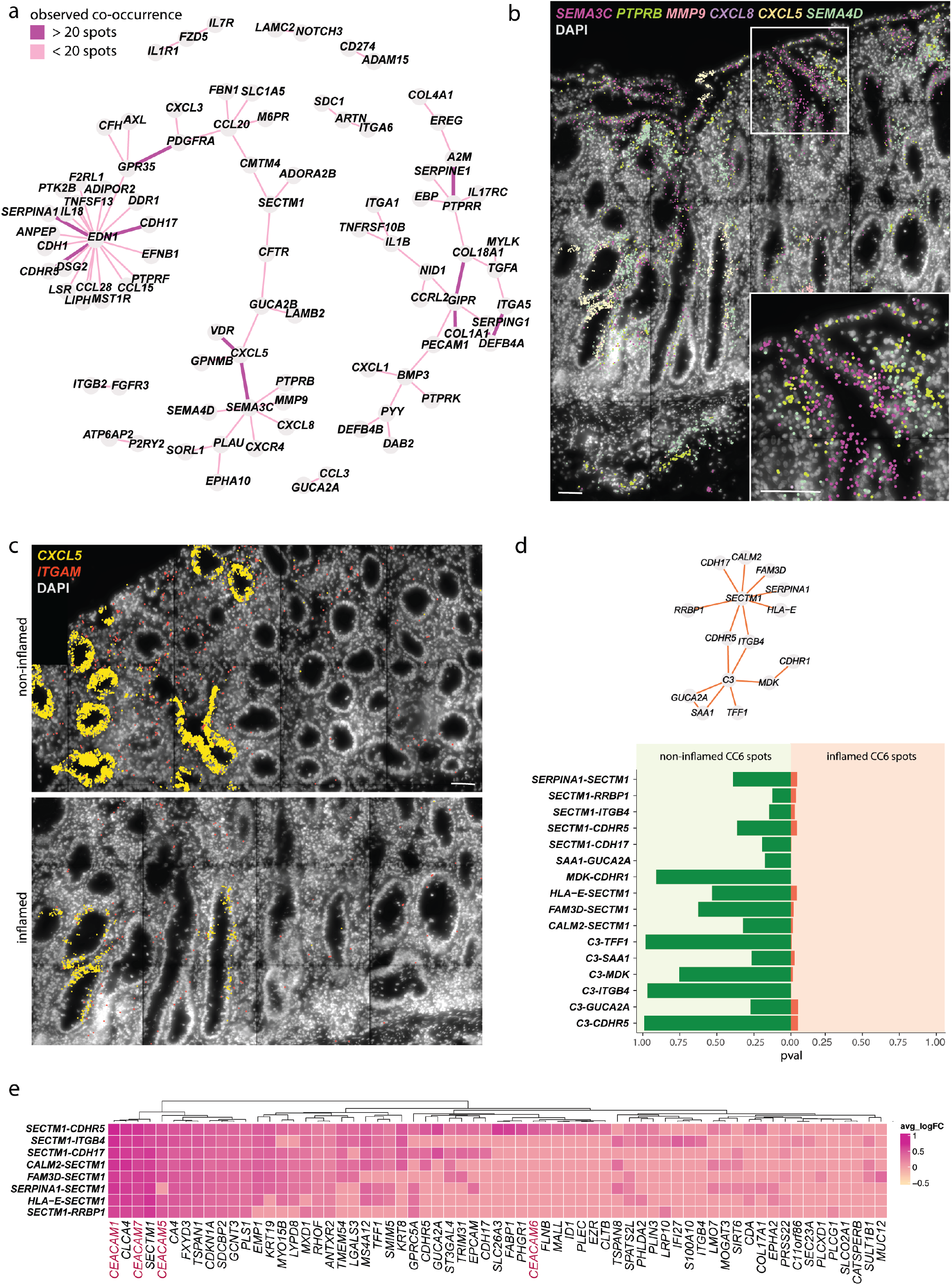
Differential co-occurrence identifies niche or condition-specific interactomes. **a,** Network representation of the CC5-specific differentially co-occurring interactome. Scale bar, 20 μm. **b,** SEMA3C co-occurrence network visualized by Molecular Cartography. **c,** CXCL5 and ITGAM expression in non-inflamed and inflamed colon samples. Scale bar, 20 μm. **d,** Top, network representation of differentially co-occurring interactome within inflamed CC6 spots. Bottom: P value of differentially co-occurring interactions in non-inflamed (left, green) and inflamed (right, orange) CC6 spots. **e,** Heatmap of *SECTM1*-interactions-associated DEGs.

Molecular Cartography further revealed highly spatially restricted and sporadic epithelial expression of *CXCL5* in UC colon samples, which coincided with neutrophil-rich areas, identified by *ITGAM* transcript (**Fig 4c**). CXCL5 expression is characteristic of colon adenocarcinoma^27–29^ and colitis^30–32^, but CXCL5+ cells are also occasionally found scattered in the healthy colon^31^, indicating highly localized induction of this cytokine. DEGs associated with differentially co-occurring *CXCL5*-interactions (**Supplemental Fig S3a**), indicate a strong SOX2 and HIF1-α transcriptional signature and enrichment of gene sets associated with innate immunity and neutrophil degranulation (**Supplemental Fig S3b**). Interestingly, SOX2-induced expression of CXCL5 was previously linked to neutrophil recruitment in non-small-cell lung cancer^33^. Whether this signaling axis is involved in IBD is largely unknown. Moreover, the observation of scattered *CXCL5+* crypts in both inflamed and non-inflamed samples may indicate that aberrant, localized *CXCL5* expression is an early event in UC pathogenesis, and warrants further investigation.

These data suggest that differential co-occurrence analysis of the interactome reveals concerted tissue transcriptional responses that only arise in specific cellular neighborhoods. While spatial co-occurrence does not necessarily represent physical binding nor molecular interaction between LR pairs, ligands and their cognate receptors are bound to be co-occurring in close spatial proximity within a tissue for paracrine and juxtacrine signaling. Whether differential co-occurrence analysis can be used to predict novel putative LR pairs based on their spatial distribution remains to be orthogonally validated.

### Differential co-occurrence reveals an altered crypt interactome

Inflammation may not only generate new CNs, but also alter the architecture of existing ones by changing the way the same cell types interact with each other (e.g. by means of soluble factors). Changes in the interactome would then be reflected in altered patterns of positively co-occurring LR pairs. We tested this hypothesis in the homeostatic CC6, which is present in both inflamed and non-inflamed samples and maps onto the lower colonic crypt, where colonic stem cells reside (**Fig 1e-g**). We thus computed differential interactome co-occurrence between inflamed and non-inflamed spots of CC6. 16 pair-wise LR combinations are differentially co-occurring in inflamed samples *(P_inflamed_ > 0.05* & *P_non-inflamed_ > 0.05*) (**Fig 4d**). As above, by performing this analysis across conditions but within the same CC, we filtered out effects arising from different cell type compositions, and recovered the differential interactions of the same groups of cells. Specifically, we detected differential co-occurrence of *Complement 3* (C3) with *MDK* and *SAA1*, all of which have roles in the recruitment of leukocytes^34–37^. In addition, inflammation-specific differential co-occurrence of *C3* and trefoil factor 1 *(TFF1)* might indicate a tissue-protective program, as both factors were implicated in epithelial maintenance^38,39^.

The differentially co-occurring LR network centered around *SECTM1* (Secreted And Transmembrane 1), similarly suggests a balance between immune cell recruitment and tissue protection. SECMT1 is a soluble protein highly expressed by crypt-top colonocytes (mostly in CC1), that exerts immunomodulatory and chemotactic functions via CD7 on T cells^40–42^. In CC6 (crypt bottom), we observed its expression mostly in DCs *(CLEC10A+)* and neutrophils *(ITGAM+)* (**Supplemental Fig S3c**) but it’s expression was also recently reported in intestinal epithelial stem cells^43^. In inflamed samples, *SECTM1* differentially co-occurs with *FAM3D* - a monocyte and neutrophil^44^ chemoattractant secreted by TAs and Goblet cells, promoting homeostasis during colitis^45^ - and *SERPINA1*, a Goblet-cell derived tissue-protective factor and inhibitor of neutrophil-derived elastase^46^ **(Fig 4d)**. Interestingly, among DEGs associated with *SECTM1*- interactions are several *CEACAM* genes, encoding for epithelial-derived adhesion proteins involved in immune modulation and colitis^47,48^ **(Fig 4e)**. Collectively, *SECTM1*-interactions-associated DEGs are enriched in genes involved in the assembly of hemidesmosomes, whose integrity protects the epithelium against inflammation^49^ (**Supplemental Fig S3d**). These data suggests that in inflamed colon areas, the crypt epithelium collectively responds to insults by secreting tissue protective factors and by enhancing cell adhesion. Interestingly, *SECTM1* is one of the genes that is differentially expressed between pediatric responders and nonresponders to anti-TNF therapy for IBD^50^. However, while the immuno-modulatory function of this protein in cancer has been the focus of several studies^40,42,51^, its role in colitis is largely unknown.

### Repair pathways are conserved across species

Finally, we applied co-occurrence analysis to an independent Visium dataset of the healing mouse colon^52^ (**Fig 5a**). As no matched scRNASeq dataset was available, we deconvoluted spots in a reference-free manner using topic modeling (see Methods), and clustered the deconvoluted matrix in 8 CCs (**Fig 5b**).

**Figure 5.**
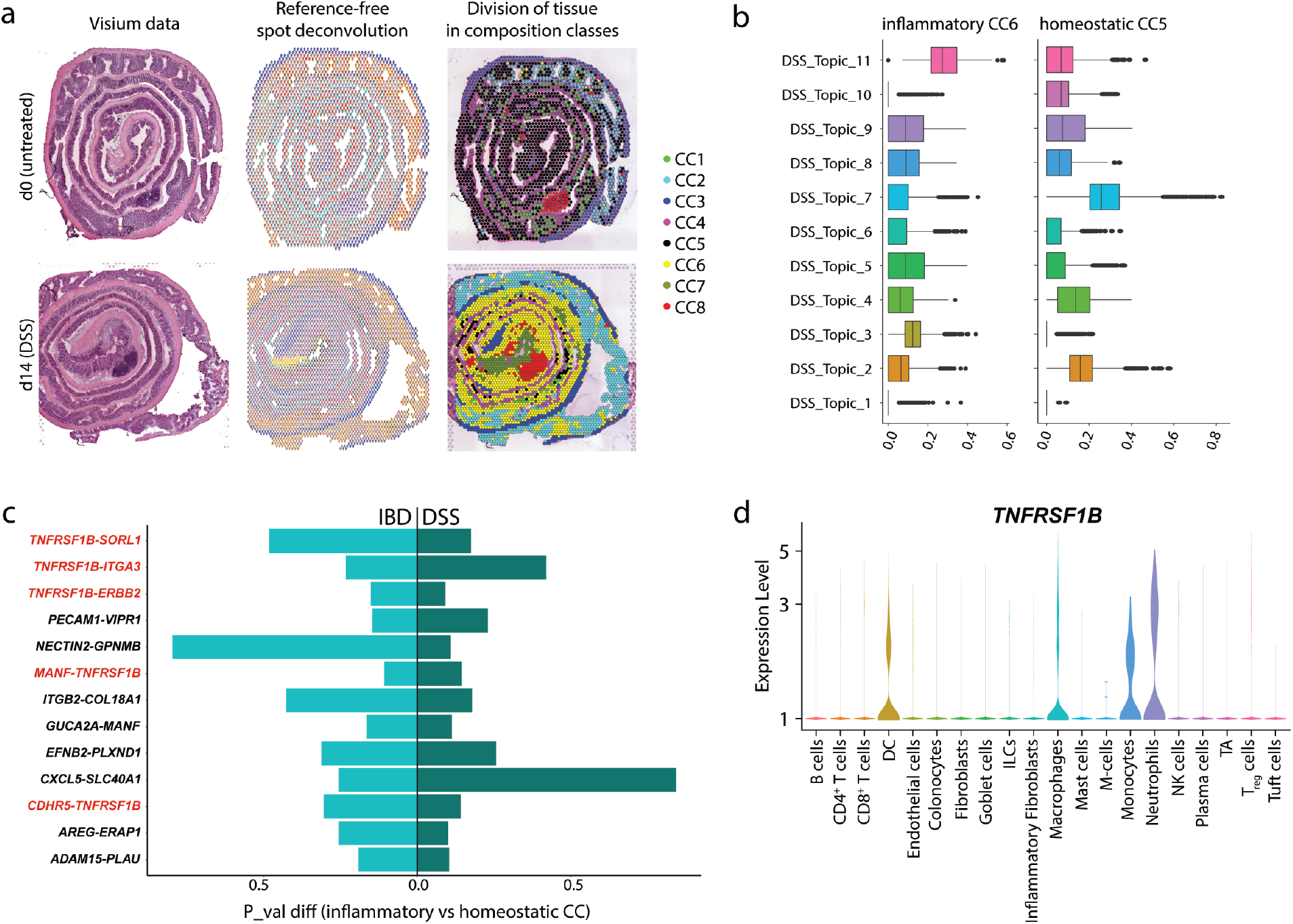
ISCHIA identifies conserved co-occurring LR pairs. **a,** Visium data from Parigi *et al*, 2022^52^, representing untreated (d0) and regenerating (d14 post DSS) mouse colon. Left: hematoxylin-eosin staining, center: reference-free spot deconvolution, right: division of tissue in composition classes. **b,** Topic distribution in inflammatory and homeostatic CCs in mouse colon. **c,** Common differentially co-occurring LR pairs with *P_inflammatory_CC_ < 0.05 & P_homeostatic_CC_ > 0.05* in both human IBD (left) and mouse DSS (right) Visium data. **d,** Expression of *TNFRSF1B* in human IBD scRNASeq reference.

Next, in both the human and murine dataset, we identified LR pairs that are differentially co-occurring in inflammatory vs homeostatic CCs (P_inflammatory_CCs_ < 0.05 & P_homeostatic_CCs_ > 0.05). Interestingly, we found several orthologous LR pairs that were differentially co-occurring in both mouse and human inflammatory CCs, and whose molecular interaction is also reported in PPI databases (**Fig 5c**). Several of those pairs involve *TNFRSF1B*, encoding for the tumor necrosis factor receptor 2, and a genetic risk factor for colitis. Indeed, polymorphisms in *TNFRSF1B* are associated with increased susceptibility to IBD^53–55^. However, little is known about the function of this gene in inflammation^56^. Thanks to our comprehensive IBD scRNAseq reference, we could map the expression of *TNFRSF1B* in humans to DCs, monocytes, macrophages and, most prominently, neutrophils (**Fig 5d**). Recently, TLR2 signaling in colitis was linked to increased extracellular trap formation by neutrophils^57^, which are emerging as key cellular players IBD^58^. However, to what extent this pathway contributes to tissue damage during IBD is still unknown, and highlights the need for cellular atlases that include granulocytes.

Collectively, our comparative analysis in human and mouse colitis suggests that some repair CNs may be shared across species, and illustrates the ability of ISCHIA to identify co-occurring LR pairs from ST data also in the absence of a single cell reference.

## Discussion and conclusions

In community ecology, co-occurrence analysis is the study of interactions between distributions of species at defined spatial locations^6^. While the link between co-occurrence and ecological interactions is still debated, modeling of presence-absence data has been instrumental in understanding the rules of ecological community assembly^59^. Here, we propose the use of co-occurrence for the analysis of spatial transcriptomics data. We exploit the fact that proximity is a prerequisite for juxtacrine and paracrine cell-cell communication, which in turn constitutes the basis for the coordinated function of CNs. We reasoned that spot data from sequencing-based ST methods such as Visium is ideal for co-occurrence analysis, as it simultaneously captures - at multiple spatially restricted locations - information about 1) neighboring cell types, 2) expressed LR genes, and 3) associated transcriptional responses.

We thus generated Visium data from human UC colon samples, deconvoluted spot data based on a comprehensive scRNASeq IBD reference, and grouped spots in composition classes. Composition-based clustering of the tissue represents a major advantage of this method. Indeed, the downstream cell interaction predictions are conducted within individual CCs, filtering out transcriptomic differences arising from distinct cell type compositions. We then performed co-occurrence analysis of cell types within CCs. Next, we linked co-occurrence of cell types to co-occurrence of LR pairs, reconstructing nodes and edges of CNs. Finally, we integrated cell types, LR genes and associated gene signatures to infer CN function. Thereby, we identify a M-cell-fibroblast network in the inflamed colon of UC patients. We validated our findings from bulk spot data at single-cell resolution by means of Molecular Cartography. We showed that co-occurrence analysis can successfully be applied to such data, and that it can further restrict predictions from Visium data by analyzing very small cellular networks of k=5 neighboring cells.

We propose differential co-occurrence analysis of the interactome as a predictive tool to infer concerted and spatially localized tissue responses. These can be envisioned as ‘conversations’ between cells in a CN, which may vary according to the composition of the CN or the environmental condition the CN is in (inflamed vs non-inflamed colon). We define differential co-occurrence as a condition- or niche-specific positive co-occurrence between interactome components *(P_condition_1_ < 0.05 & P_condition_2_ > 0.05*). Thereby, we identify differentially co-occurring LRs specific to the inflammatory CC5 centered around *EDN1, SEMA3C*, and *CXCL5*. We further identify inflammation-induced, protective responses from the colonic crypt involving the complement cascade and the immuno-modulator *SECTM1*. Finally, we apply co-occurrence analysis to an independent mouse Visium dataset and uncover differentially co-occurring LR pairs shared between the inflamed human and murine colon.

Collectively, our data show that co-occurrence analysis can be applied to both sequencing- and imaging-based ST data, to refine interaction predictions from scRNAseq based analysis. It is complementary to differential gene expression, as it does not depend on the abundance of a given cell type or the expression levels of the analyzed genes, but rather on their spatial arrangement and distribution in the tissue. Whether spatial proximity and differential co-occurrence of LR genes can be employed, in conjunction with protein-protein interaction predictions, for the identification of novel putative interacting pairs, warrants further investigation. Nonetheless, we show that co-occurrence analysis on spatial transcriptomics data can be used to chart the distribution and infer the interactions of cell types and transcripts in the tissue, revealing disease-specific cellular communities, and predicting juxtacrine and paracrine signaling. It therefore represents a powerful tool for hypothesis-generation from spatial transcriptomics data.

## Methods

### Data

#### scRNASeq

scRNASeq data from IBD patients were obtained from published datasets^7–9^ or collected in house with 10X Genomics (2 UC patients, 2 suspected IBD patients, 4 healthy controls).

#### 10X Visium

Fresh frozen UC colon samples were sectioned onto 10X Visium Spatial Gene expression slide. cDNA libraries were generated according to the manufacturer’s instructions. After methanol fixation, tissue morphology was assessed by hematoxylin and eosin (H&E) staining. The permeabilization time of 50 min was assessed with the Tissue Optimisation Protocol. Lysis, reverse transcription, second strand synthesis and cDNA denaturation were performed on the slide.

cDNA was then amplified by PCR using the cycle number identified by qPCR and then subjected to end repair, A-tailing, adapter ligation and indexing to generate sequencing libraries. Quality and quantity of all libraries were assessed using the dsDNA high-sensitivity (HS) kit (Life Technologies #Q32854) on a Qubit 4 fluorometer (Thermo Fisher) and high sensitivity D1000 reagents and tapes (Agilent #5067-5585, #5067-5584) on a TapeStation 4200 system (Agilent Technologies). Paired-end sequencing was performed on a NovaSeq 6000 system (Illumina) using NovaSeq SP Reagent Kits (100 cycles) v1.5. Data was pre-processed using Space Ranger (v1.2.0) (10X Genomics) with GRCm38 v2020-A genecode. Visium data of DSS-treated murine colon was obtained from Parigi et al.^52^

#### Molecular Cartography

Fresh frozen UC colon samples were sectioned onto coverslips and processed by Resolve Biosciences. Cellpose^60^ (v. 2.0.4) was used to segment nuclei in the DAPI images with the pretrained nuclei model and flow_threshold 0.5, cellprob_threshold - 0.2. Using the ‘expand_labels’ function in scikit-image, the nuclear segments were then expanded by 10 pixels (1.38 μm) and transcripts were subsequently assigned to the expanded segments. Segments with less than 3 molecules or 3 genes detected were removed from the analysis.

### Integration and annotation of scRNASeq data

For every scRNASeq dataset, we performed log normalization with scale factor 10000. The top 2000 variable genes are selected for each dataset using the “vst” method in Seurat^61^. Next, we used the *FindIntegrationAnchors* function to align shared cellular populations across datasets by finding pairs of cells that are in matching biological states. The identified anchors are then used in the *IntegrateData* function to calculate an integrated (batch-corrected) expression matrix for all cells, enabling joint analysis. We used the labels provided by Smillie *et al.^7^* as reference for automatic annotation of all the clusters in the integrated data with the *TransferData* function from Seurat. W then manually grouped the clusters into 20 major cell types (Supplemental Fig. S1A). Granulocytes were directly retrieved from the Hander *et al* dataset^9^.

### Deconvolution of Visium spots

#### Reference-based deconvolution

we used the integrated scRNASeq reference to deconvolute human IBD Visium spot data with SPOTlight^62^, a computational tool that enables the integration of ST and scRNA-seq data using a seeded non-negative matrix factorization (NMF) regression. The deconvolution function was run with default parameters.

#### Reference-free deconvolution

Mouse colitis Visium data was deconvoluted in a reference free manner with STdeconvolute^63^, which builds on latent Drichlet allocation (LDA). Given a count matrix of gene expression in multicellular Visium spots, STdeconvolute applies LDA to infer the putative transcriptional profile for each cell type and the proportional representation of each cell type in each multi-cellular spot. The deconvolution function was run with default parameters.

### Co-occurrence analysis on Visium data

ISCHIA uses spatial co-occurrence, a probabilistic approach inspired by species co-occurrence models in ecology that assigns a measurable property for spatial proximity between cell types or interactome component genes. Observed co-occurrence is quantified as the number of spatial spots where two cell types or genes co-occur. Observed co-occurrence is compared to the expected co-occurrence, where the latter is the product of the two cell types’ probability of occurrence, multiplied by the number of spatial spots:

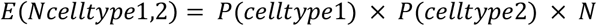

This probabilistic model uses combinatorics to determine whether the observed frequency of co-occurrence is significantly greater than expected (positive association), significantly smaller than expected (negative association), or not significantly different to expected (random association). Specifically, this analysis calculates, for each cell type pair, the exact probability of the observed co-occurrence to be greater than (P(gt)) or less than (P(lt)) the expected co-occurrence. Importantly, this analysis is distribution-free and the results can be interpreted and reported as p-values, without reference to a statistic. Therefore, given two cell types in a dataset, a P(lt) ≤ α suggests that those two species are negatively associated (where P(lt) = $p_lt and α = 0.05).

To define the cell type composition of each spot, the spot transcriptional profile is deconvoluted. The resulting spot × cell type/topic probability matrix is then converted to a binary presence-absence matrix by thresholding (cell type deconvoluted probability > 0.1 = presence). For L-R pairs, the presence-absence matrix is derived by their expression. This matrix is then used as an input for co-occurrence calculation with the R package cooccur^64^: we calculate the probability of selecting a spot that has cell type #1 given that it already has cell type #2.

The probability that the two cell types co-occur at exactly j number of spots is given by

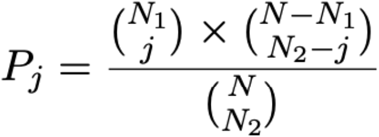

For *j* = 1 to *N*_1_ spots:

*N*_1_ = number of spots where cell type #1 occurs

*N*_2_ = number of spots where cell type #2 occurs

*N* = total number of spots sampled (where both cell types could occur).

The term, 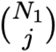 represents the number of ways of selecting *j* spots that have cell type #1 given that there are *N*_1_ such spots in the “population” of all spots.

The term 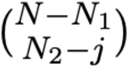 represents the number of ways of selecting *N*_2_ – *j* spots that have cell type #2 but not cell type #1 given that there are *N* – *N*_1_ such spots.

The numerator 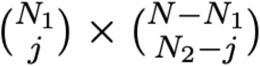 gives the total number of ways of selecting *j* spots that have cell types #1 and #2. The denominator 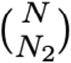 represents the total number of ways that *N*_2_ number of spots could be obtained out of a total of *N* spots. Thus the equation is giving the proportion of the *N*_2_ spots that also have cell types #1 under the condition that the two cell types co-occur at *j* spots.

The spatial co-occurrence for L-R pairs follows the same principle (presence-absence matrix based on count > 1). Additionally, we rank L-R pairs by calculating the correlation of expression between the ligand and receptor gene in each spatial spot, expecting that if a L-R pair is interacting, the expression of the ligand and receptor should correlate as well.

### Co-occurrence analysis on Molecular Cartography data

We applied co-occurrence analysis on Molecular Cartography data by transforming the x,y coordinates of the segmented cells into cellular neighborhoods. To do so, we calculated the K-nearest neighbor graph of the x,y coordinates where k=5. We applied ISCHIA’s co-occurrence analysis on the presence-absence matrix of calculated CNs and cell types as explained above.

### Graphical illustrations

Schematics of experimental workflows were created using a licenced version of Biorender.com.

## Supplemental Information

- Additional file 1: Supplemental figures 1–3.
- Additional file 2: Supplemental Table 1.

## Data access

The datasets supporting the conclusions of this article are available in the Zenodo repository, [10.5281/zenodo.7589581]. The code used in this study is available at: <u>ati-Iz IISCHIA: Framework for analysis of cell-types and Ligand-Receptor cooccurrences</u> (github.com)

## Acknowledgements

We thank members of the Moor lab and the Roche Immunology Discovery department for insightful discussions. We also thank D. Eletto for organizing our lab retreat on the island of Ischia, where this project was conceived.

## Author contributions

A.L. and J.K. performed the experiments. A.L., C.B. and K.B. performed the analysis. A.L., C.B. and X.F. wrote the manuscript. A.E.M and A.F. supervised the study.

---

**Supplemental Figure S1.**
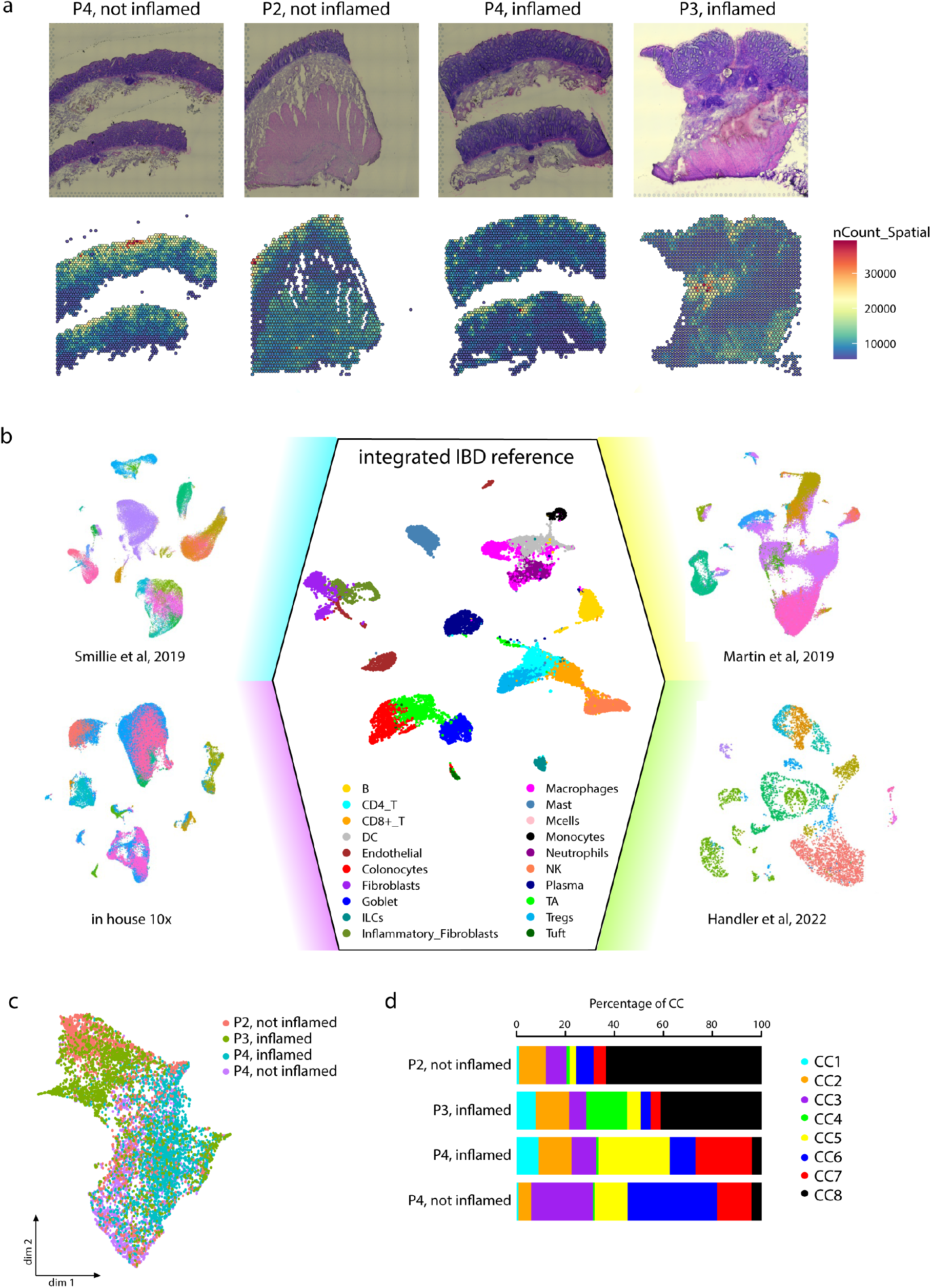
Composition-aware clustering of human colon Visium data. **a,** Top: Hematoxylin eosin staining of colon resections from four patient samples (a total of 3 patients, 2 inflamed and 2 not inflamed samples) analyzed by Visium ST (10X Genomics). Bottom: Capture area colored by number of sequenced UMI. **b,** Integration of published and in house IBD scRNASeq datasets yields a comprehensive IBD scRNASeq reference. Dots represent single cells, colored by cell type. **c,** Dimensionality reduction of spots colored by patient. **d,** Percentage of CC per patient.

**Supplemental Fig S2.**
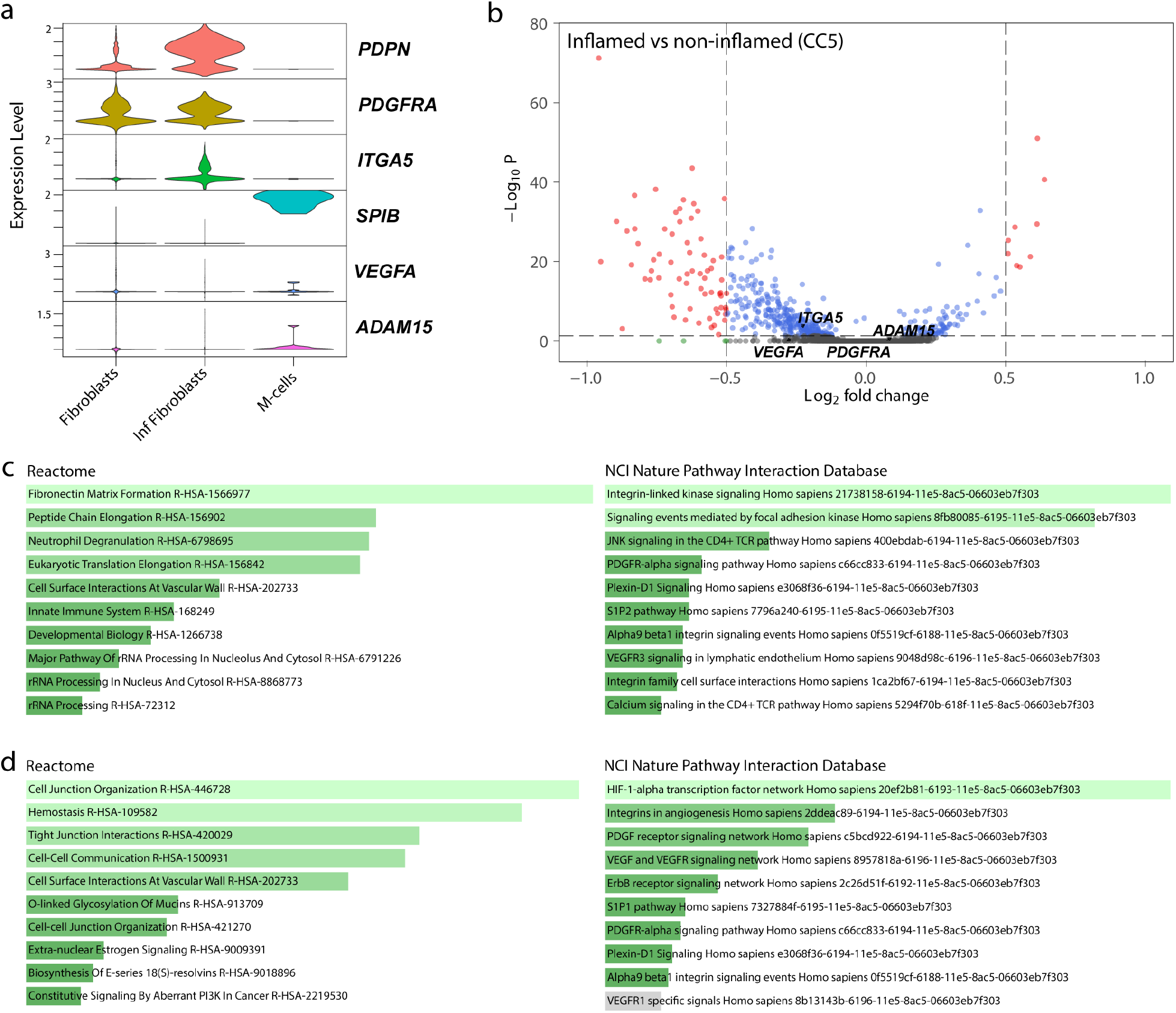
Co-occurring LR pairs are associated with a transcriptional signature. **a,** Expression of cell type markers *(PDPN, PDGFRA, SPIB)* and LR pairs *(PDGFRA, VEGFA, ITGA5, ADAM15)* by fibroblasts and M-cells in integrated IBD scRNASeq reference. **b,** Differential gene expression within CC5, inflamed vs non-inflamed (4 samples, 3 patients). **c**, Reactome and NCI Nature Interaction Database pathway enrichment of *ITGA5-ADAM15*-associated genes. **d**, Reactome and NCI Nature Interaction Database pathway enrichment of *VEGFA-PDGFRA*-associated genes.

**Supplemental Fig S3.**
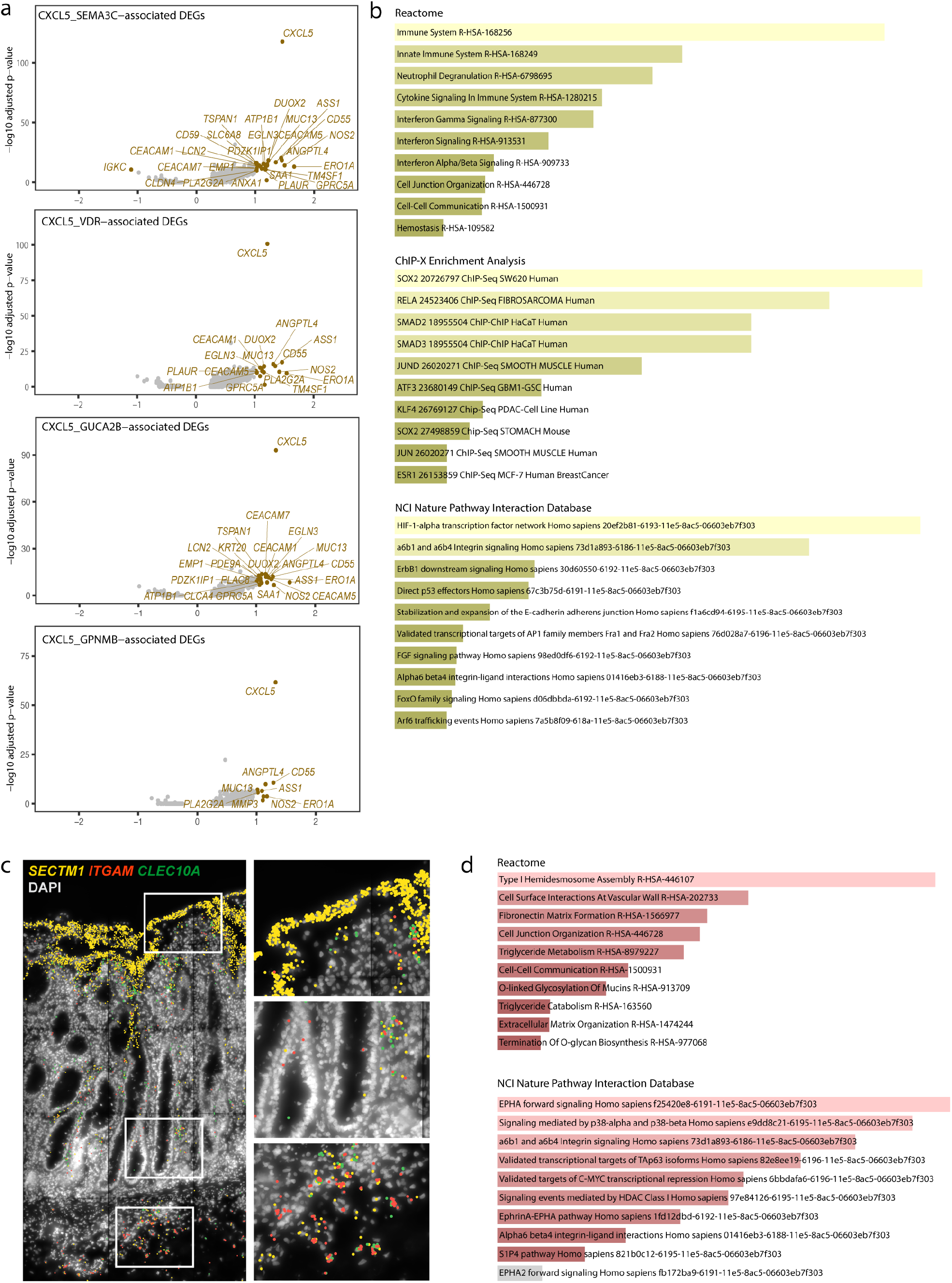
Differential co-occurrence analysis of the interactome reveals concerted tissue responses. **a,** *CXCL5*-interactions associated DEGs. **b,** Reactome, ChiP-X Enrichment Analysis andNCI Nature pathway enrichment of shared *CXCL5*-associated DEGs. **c,** Molecular Cartography image of *SECTM1, ITGAM* (neutrophils) and *CLEC10A* (DCs) expression. **d**, Reactome and NCI Nature pathway enrichment of *SECTM1*-associated DEGs.

